# EigenGWAS: finding loci under selection through genome-wide association studies of eigenvectors in structured populations

**DOI:** 10.1101/023457

**Authors:** Guo-Bo Chen, Sang Hong Lee, Zhi-Xiang Zhu, Beben Benyamin, Matthew R. Robinson

## Abstract

We apply the statistical framework for genome-wide association studies (GWAS) to eigenvector decomposition (EigenGWAS), which is commonly used in population genetics to characterise the structure of genetic data. The approach does not require discrete sub-populations and thus it can be utilized in any genetic data where the underlying population structure is unknown, or where the interest is assessing divergence along a gradient. Through theory and simulation study we show that our approach can identify regions under selection along gradients of ancestry. In real data, we confirm this by demonstrating *LCT* to be under selection between HapMap CEU-TSI cohorts, and validated this selection signal across European countries in the POPRES samples. *HERC2* was also found to be differentiated between both the CEU-TSI cohort and within the POPRES sample, reflecting the likely anthropological differences in skin and hair colour between northern and southern European populations. Controlling for population stratification is of great importance in any quantitative genetic study and our approach also provides a simple, fast, and accurate way of predicting principal components in independent samples. With ever increasing sample sizes across many fields, this approach is likely to be greatly utilized to gain individual-level eigenvectors avoiding the computational challenges associated with conducting singular value decomposition in large datasets. We have developed freely available software to facilitate the application of the methods.

## Introduction

In population genetics, eigenvectors have been routinely used to quantify genetic differentiation across populations and to infer demographic history (Cavalli-Sforza *et al.*, 1996; Novembre *et al.*, 2008; Reich *et al.*, 2009). More recently, eigenvectors are commonly used as covariates in genome-wide association studies (GWAS) to adjust for population stratification (Price *et al*., 2006). Eigenvectors are usually estimated for each individual (individual-level eigenvectors, involving the inversion of a *N* × *N* matrix, where *N* is sample size). Theoretical studies have suggested that individual-level primary eigenvectors are measures of population differentiation reflecting *F_st_* among subpopulations (Patterson *et al*., 2006; McVean, 2009; Bryc *et al.*, 2013) and can be interpreted as the divergence of individuals from their most recent common ancestor. Eigenvectors can also be estimated for each SNP (SNP-level eigenvectors, which involve inversion of a matrix, is the number of SNPs) and these SNP-level eigenvectors can be interpreted as *F_st_* metrics of each SNP (Weir, 1996). SNP-level eigenvectors from a reference population are useful for revealing the population structure of independent samples (Zhu *et al*., 2008) as they can be used to project, or predict, the eigenvector values of individuals. However, due to high-dimensional nature of GWAS data (commonly expressed as *M* ≫ *N*), direct estimation of SNP-level eigenvectors is nearly impossible when using millions of single nucleotide polymorphisms (SNPs).

Singular value decomposition (SVD) enables SNP-level eigenvalues to be obtained in a computationally efficient manner for any set of genotype data (Chen *et al*., 2013), however, it is not possible to determine the SNPs that contribute most to the leading eigenvector, or to test whether specific SNPs are differentiated along the genetic gradient described by the eigenvector. Here, we propose an alternative simple, fast approach for the estimation of SNP-level eigenvectors. By using individual-level eigenvectors as phenotypes in a linear regression, we demonstrate that the regression coefficients generated by single-SNP regression are equivalent to SVD SNP effects as proposed by Chen et al (Chen *et al*., 2013). As the single-SNP regression resembles the popular single-marker GWAS method, as implemented in PLINK (Purcell *et al*., 2007), we call this method EigenGWAS. We show that the EigenGWAS framework represents an alternative way for identifying regions under selection along gradients of ancestry.

## Results

### Properties of the estimating SNP effects for eigenvectors

We applied EigenGWAS to the HapMap cohort, a known structured population. Eigenvectors were estimated via principal component analysis based on the ***A*** matrix using all 919,133 SNPs. We conducted EigenGWAS for HapMap, using *E_k_*, the *k^th^* eigenvector, as the phenotype and investigated the performance of EigenGWAS from *E*_1_ to *E*_10_. From *E*_1_ to *E*_10_, we found 546,716 significant signals (231,677 quasi-independent signals after clumping) on *E*_1_ and gradually reduced to 236 (163 after clumping) selection signals on *E*_10_ (**Fig. 1**). The large number of genome-wide significant loci are likely because HapMap3 was comprised of samples from different ethnicities, and these loci can be interpreted as ancestry informative marker (AIM). For each *E_k_,* its associated eigenvalue was highly correlated with the *λ_GC_,* the genomic inflation factor that is commonly used in adjusting population stratification for GWAS (Devlin and Roeder, 1999), resulted from its EigenGWAS. The top five eigenvalues associated to HapMap samples were 100.14, 47.66, 7.168, 5.92, and 4.40, and the corresponding *λ_GC_* of EigenGWAS were 103.72, 44.69, 6.47, 5.17, and 3.96, respectively (**Table 1**). The large eigenvalues observed were consistent with previous theory that the magnitude of eigenvalues indicating structured population (Patterson *et al*., 2006). The connection between *λ_GC_* and eigenvalues, provides a straightforward interpretation: a large *λ_GC_* indicates underlying population structure (Devlin and Roeder, 1999). Therefore, correction for will filter out signals due to population stratification, allowing loci under selection to be identified. These observations agreed well with our theory (see Methods & Materials).

**Figure 1.**
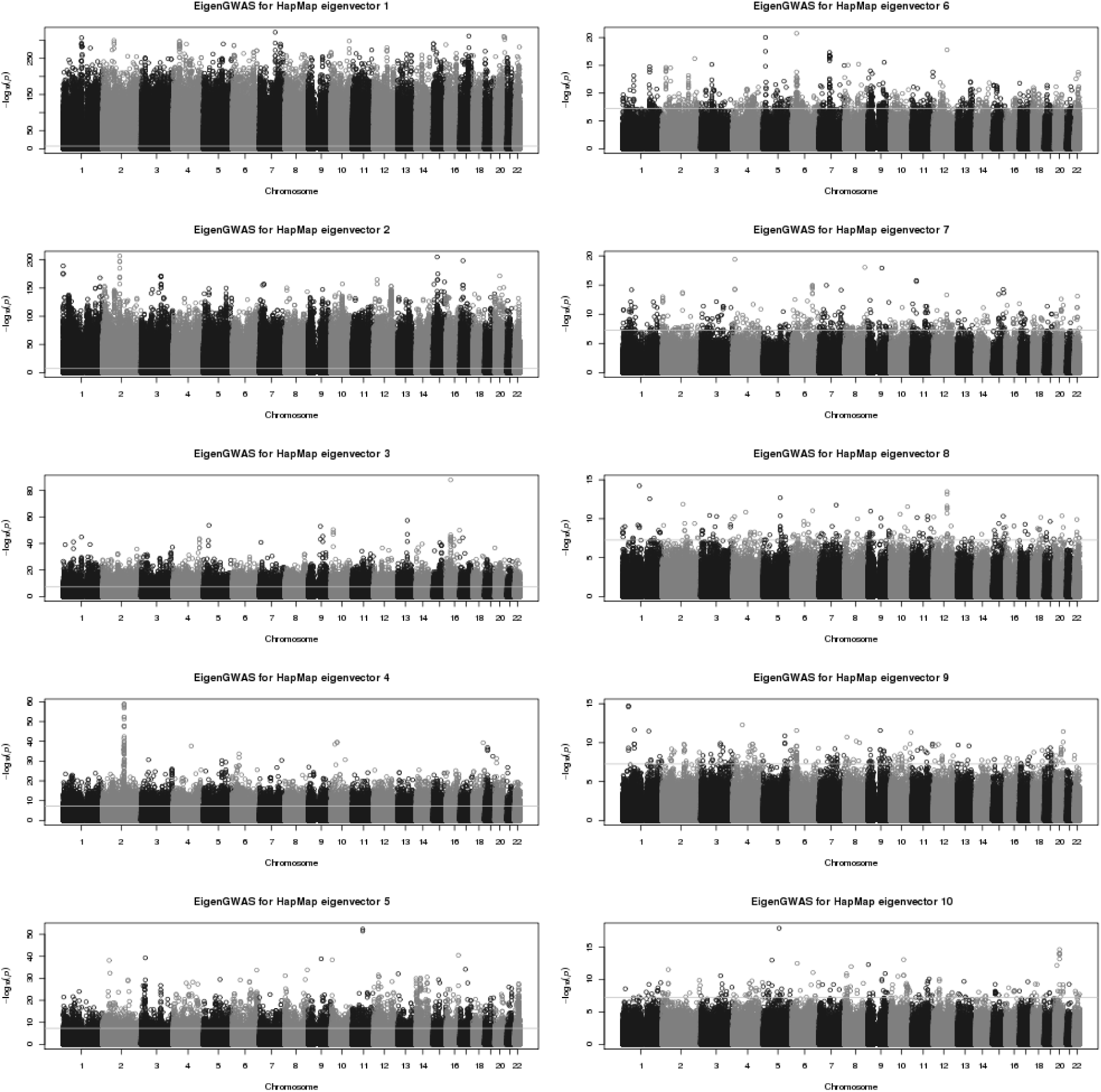
Manhattan plots for EigenGWAS for top 10 eigenvectors for HapMap. Using *E_i_* as the phenotype, the single-marker association was conducted for nearly 919,133 markers. The left panel illustrates from *E*_1_ ~ *E*_5_; the right panel from *E*_6_ ~ *E*_10_. The horizontal lines indicate genome-wide significant after Bonferroni correction.

**Table 1.**
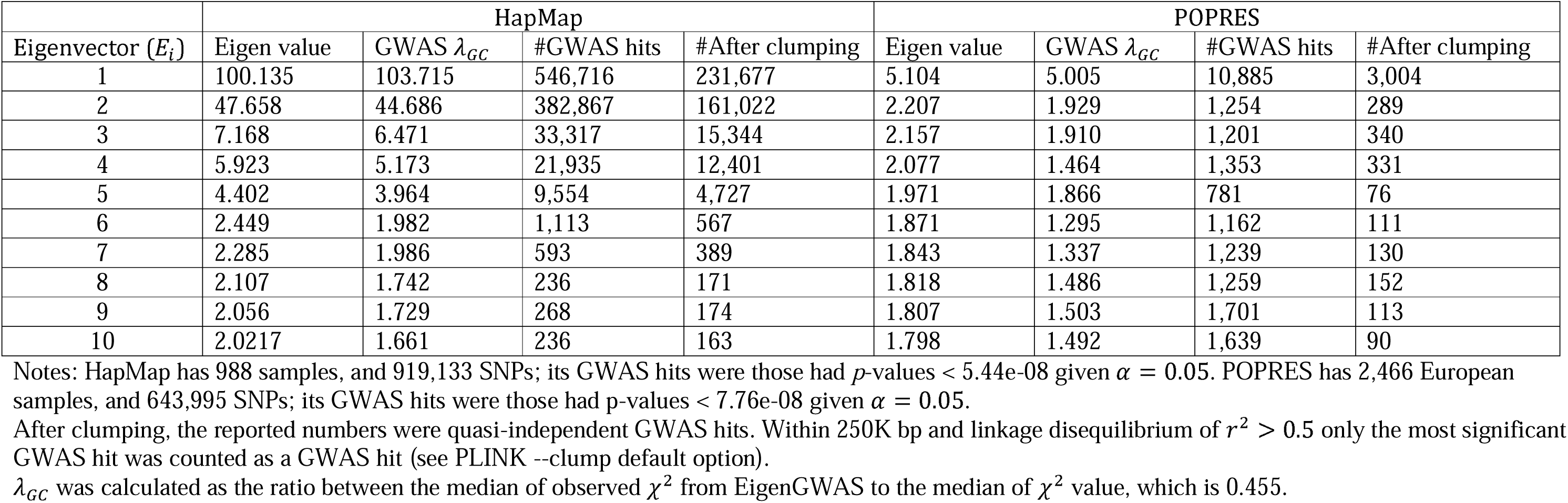
GWAS signals for on eigenvectors for HapMap and POPRES.

We demonstrate theoretically that for EigenGWAS, the estimated SNP effects using single-marker GWAS are equivalent to the estimates from BLUP, and the correlation between the estimates from these two methods was very high (greater than 0.98 on average) (**Fig. 2**), even in HapMap samples that consist of a mix of ethnicities where the ***A*** matrix is non-zero for off-diagonal elements (**Supplementary Fig. 1**). This confirms that our EigenGWAS approach provides an accurate representation of the SNP effects on eigenvalues.

**Figure 2.**
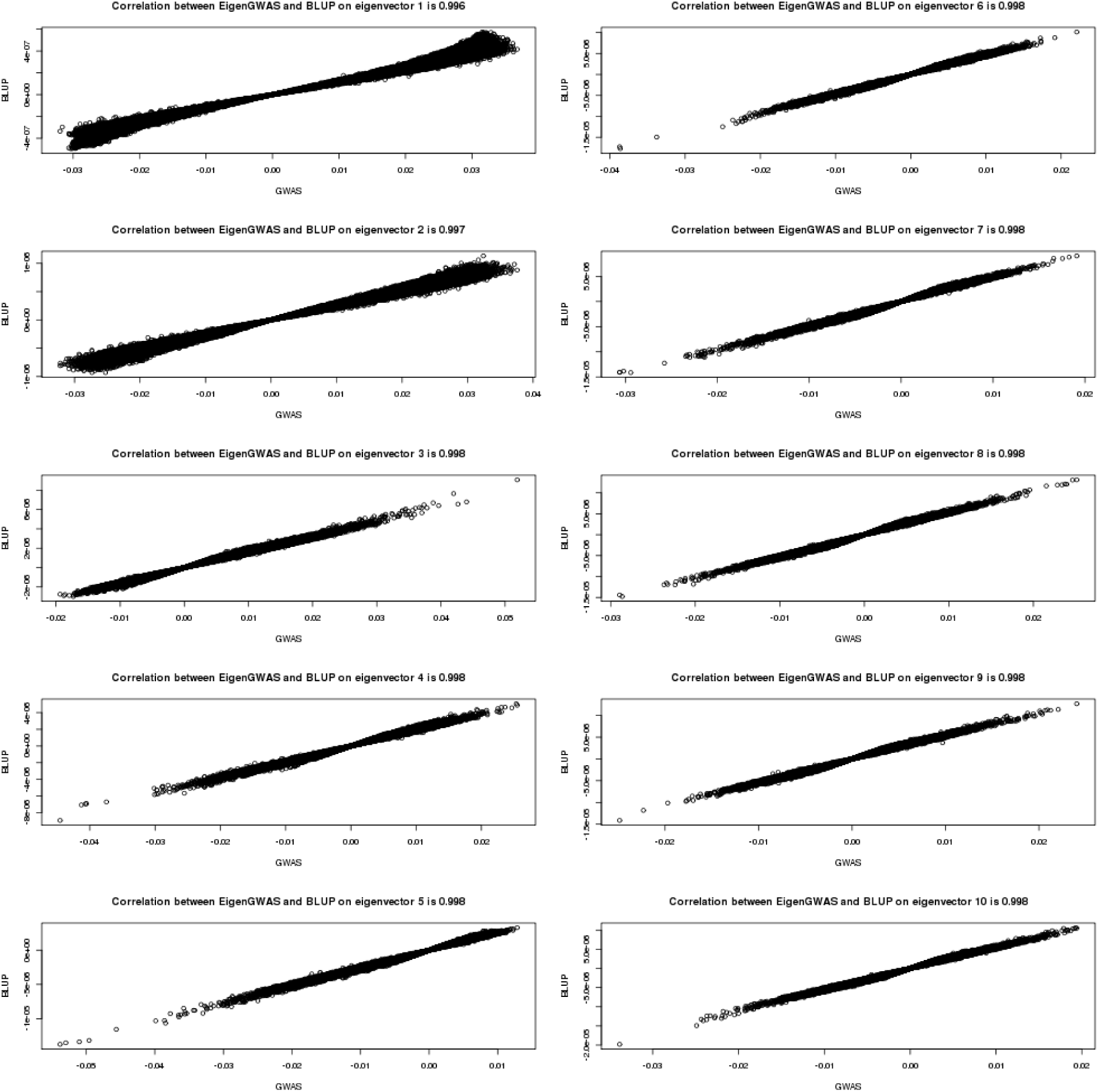
Correlation for the SNP effects estimated using EigenGWAS and BLUP for HapMap3. The x-axis represents EigenGWAS estimation for SNP effects, and the y-axis represents BLUP estimation for SNP effects. The left panel illustrates from *E*_1_ ~ *E*_5_; the right panel from *E*_6_ ~ *E*_10_. As illustrated, the correlation is nearly 1.

We also conducted EigenGWAS on the POPRES samples, from which we selected 2,466 European samples. On *E*_1_, there were 10,885 (3,004 quasi-independent signals after clumping) genome-wide significant signals, and reduced to 1,639 (90 after clumping) on *E*_10_ (**Table 1**). As in the HapMap sample, we observed a concordance between eigenvalues and *λ_GC_* in POPRES. The top five eigenvalues were 5.104, 2.207, 2.157, 2.077, and 1.971, with their associated EigenGWAS *λ_GC_* were 5.005, 1.929, 1.910, 1.464, and 1.866, respectively (**Table 1**), indicating population structure. The genetic relationship matrix (GRM) estimated from the POPRES data resembled a diagonal matrix, which had off-diagonal elements close to zero, suggesting that POPRES is a more homogenous samples as compared to HapMap (**Supplementary Fig. 1**). Correlations between the estimates from EigenGWAS and BLUP were high, with an average of greater than 0.999 from *E*_1_ to *E*_10_ (**Supplementary Fig. 2**), close to one as expected.

The chi-square statistics of the estimated SNP effects on eigenvectors from EigenGWAS were correlated with *F_st_* for each SNP, consistent with previous established relationship between eigenvectors and *F_st_* (Patterson *et al*., 2006; McVean, 2009). Using naïve threshold of *E_k_* > 0, 2,466 POPRES samples were divided into nearly two even groups, which would be served as two subgroups in calculating *F_st_*. *E*_1_ > 0 split the POPRES samples into North and South Europe; samples from UK, Ireland, Germany, Austria, and Australia were in one group, and samples from Italy, Spain, and Portugal were in the other group; samples from Switzerland and France were nearly evenly split into two groups. *F_st_* for each SNP was consequently calculated based on these two groups. For every eigenvector until *E*_10_, we observed strong correlations between *F_st_* and the chi-square test statistics for EigenGWAS signals (**Fig. 3**), and the averaged correlation was 0.925 (S.D., 0.067). For example, the correlation was 0.89 (*p*-value<1e-16) between chi-square test statistics and *F_st_* for *E*_1_ in POPRES (**Supplementary Table 1**). This correlation is consistent with our theory, where *F_st_* has a strong linear relationship with its EigenGWAS chi-square test statistic.

**Figure 3.**
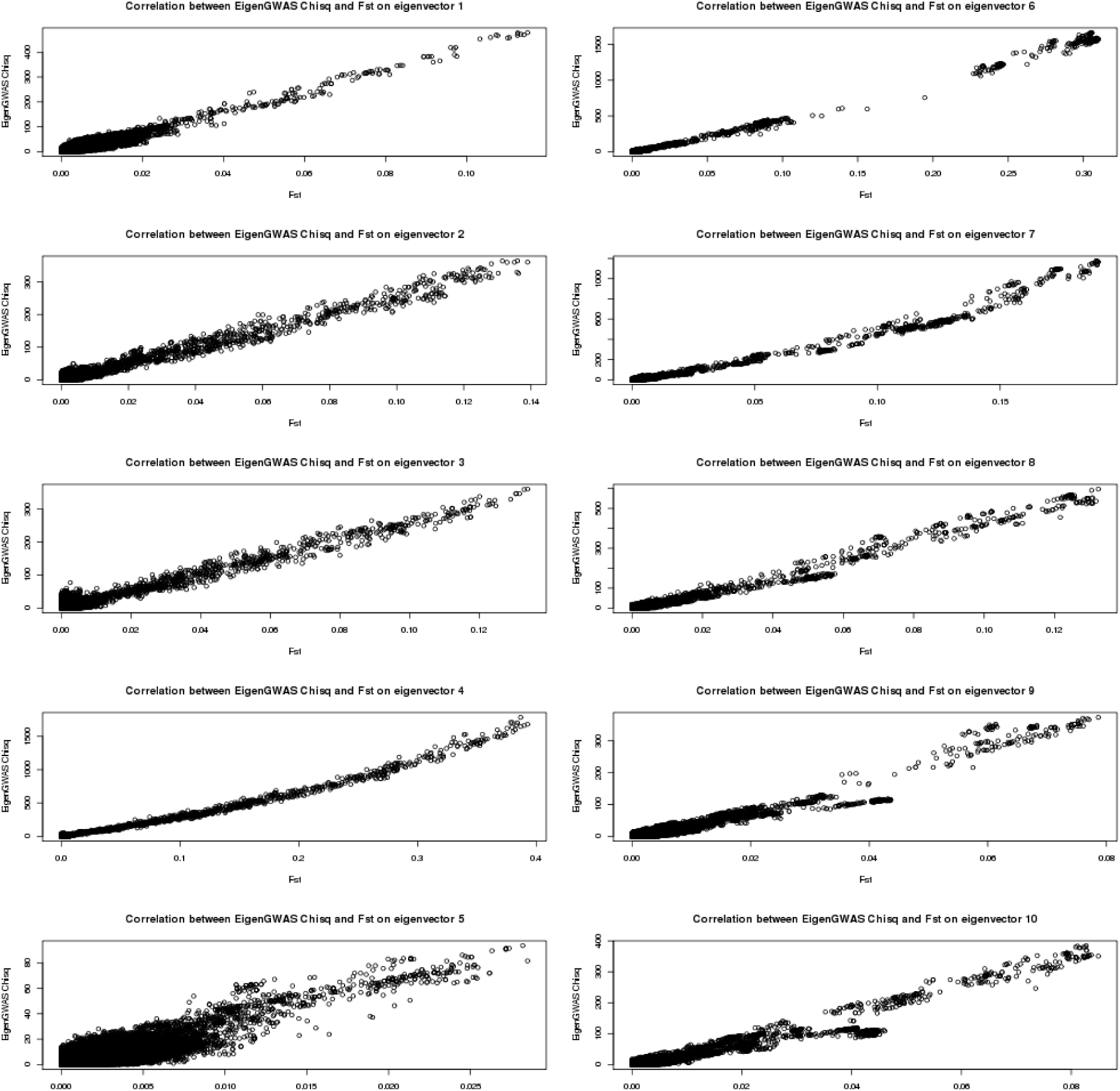
The correlation between *F_st_* and 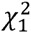 for EigenGWAS SNP effects for POPRES. For each eigenvector, upon *E_i_* > 0 or *E_i_ ≤* 0, the samples were POPRES samples were split into two groups, upon which *F_st_* was calculated for each locus.

We also validated our results in the simulation scheme I, in which there was neither selection nor population stratification. Given 2,000 simulated samples, each of which had 500,000 unlinked SNPs, the EigenGWAS showed few GWAS signals (2 genome-wide significant signals on *E*_1_, (**Supplementary Fig. 4**). After splitting the samples into 2 groups depending on *E_i_* > 0, the correlation between chi-square test statistics and *F_st_* is about 0.67 from to *E*_1_ to *E*_10_ (**Supplementary Fig. 5**). As expected, *λ_GC_* ranged from around 1.124 to 1.130, with a mean of 1.124 for EigenGWAS on the top 10 eigenvectors, indicating little population stratification for the simulated data. Furthermore, we also validated the theory in the simulation scheme II, in which there was population stratification. We wanted to know whether the adjustment of the test statistic with the greatest eigenvalue could render the distribution of the test statistics immunes of population stratification. Given various sample sizes for two subdivisions, after the adjustment for the test statistic with the largest eigenvalue, the test statistic followed the null distribution, which was a chi-square distribution of 1 degree of freedom (**Supplementary Fig. 6**), indicating a well control of population stratification after correction. The statistical power of EigenGWAS was also evaluated. As demonstrated, the power of EigenGWAS in detecting a locus under selection was determined by the ratio between the specific *F_st_* of a locus and the averaged population stratification in the sample (**Supplementary Fig. 7**).

**Figure 4.**
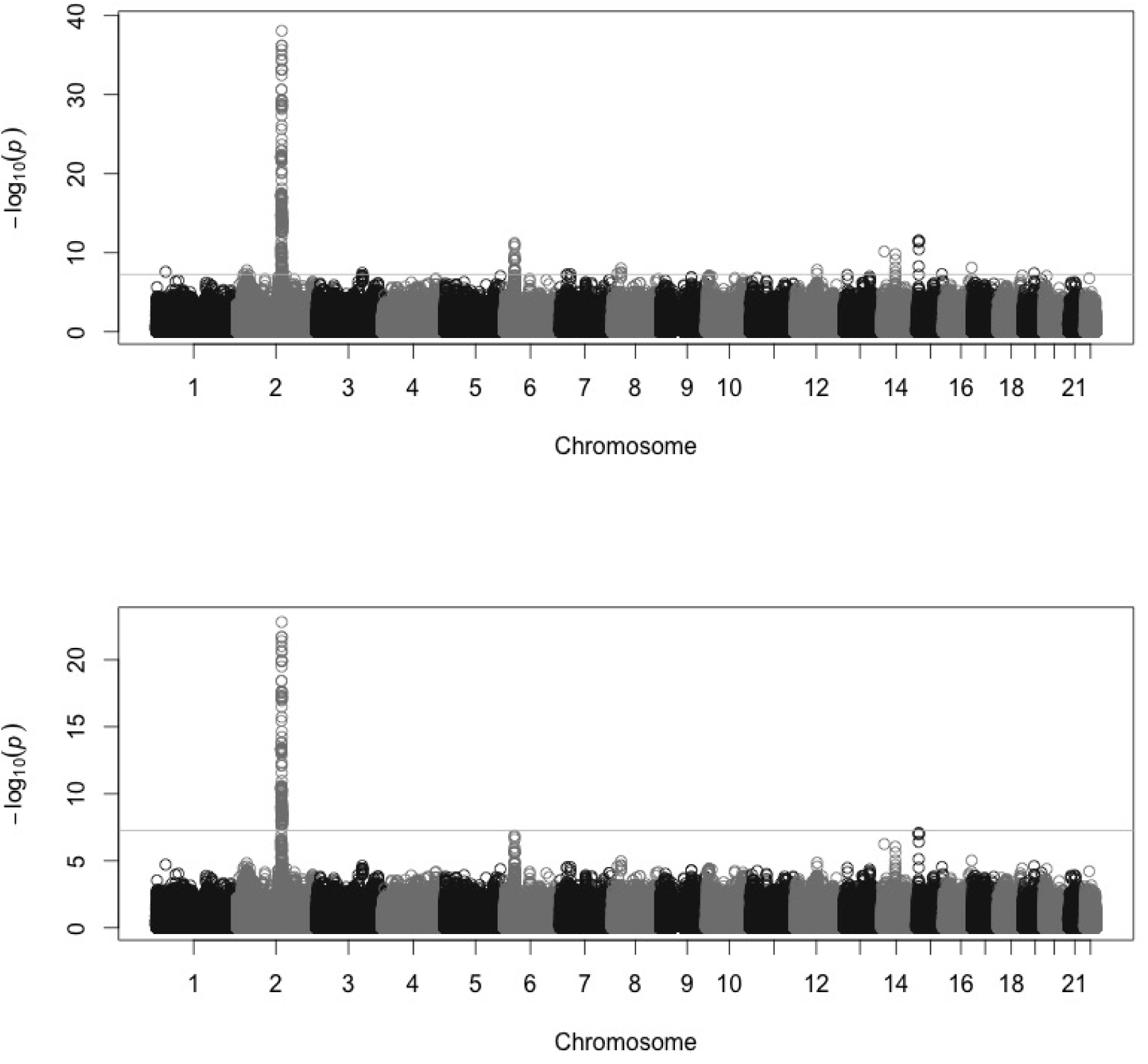
EigenGWAS for CEU (112 samples) & TSI (88 samples) from HapMap. a) Manhattan plot for EigenGWAS on *E*_1_ without correction for *λ_GC_.* When there was no correction, on chromosome 2 found *LCT,* chromosome 6 *MICA* (HMC region), chromosome 14 *HIF1A,* and chromosome 15 *HERC2.* The line in the middle was for genome-wide significant level at *α* = 0.05 given multiple correction. b) Manhattan plot for EigenGWAS on *E*_1_ with *λ*_GC_ correction, and *LCT* was still significant, and *HERC2* slightly below whole genome-wide significance level. The genome-wide significance threshold was *p*-values = 5.44e-08 for *α* = 0.05.

**Figure 5.**
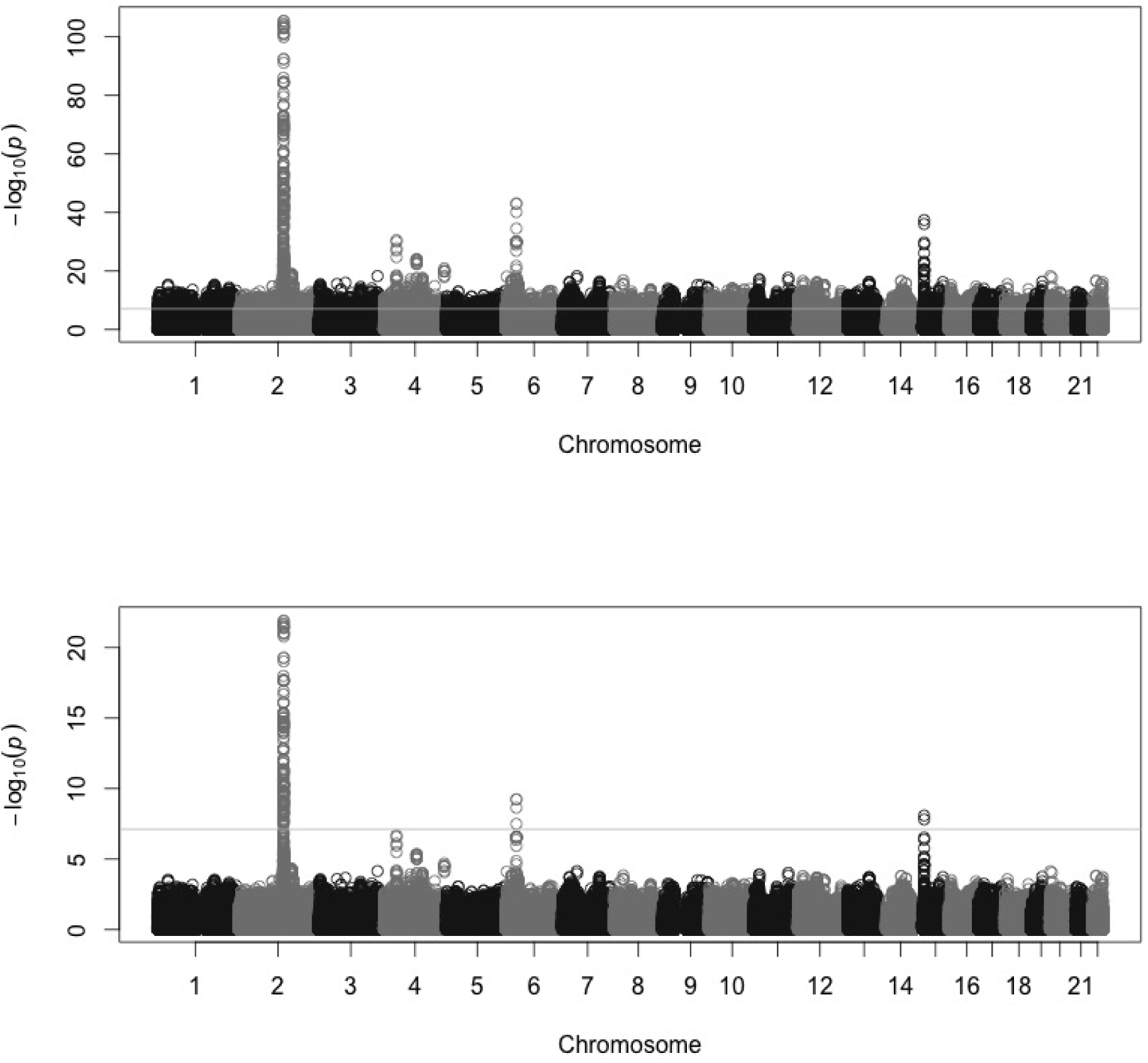
EigenGWAS for POPRES samples on eigenvector 1. a) Manhattan plot for EigenGWAS without correction for *λ_GC_.* b) After correction for λ_GC_, on Chromosome 2 found *LCT,* chromosome 6 *SLC44A4,* and chromosome 15 *HERC2.* The genome-wide significance level was *p*-values = 7.76e-08 given *α* = 0.05.

### Using EigenGWAS to identify loci under selection in structured populations

We propose EigenGWAS as a method of finding loci differentiated among populations, or across a gradiant of ancestry. Intuitively, every EigenGWAS hit is an AIM, which differ in allele frequency along an eigenvector due to genetic drift or selection. A locus under selection should be more differed across populations than genetic drift can bring out. Thus, correction for *λ_GC_,* controls for background population structure, providing a test of whether an AIM shows greater allelic differentiation than expected under the process of genetic drift.

We pooled together CEU (112 individuals) and TSI (88 individuals), which represent Northwestern and Southern European populations in HapMap. EigenGWAS was conducted on *E*_1_ > 0, which partitioned CEU and TSI into two groups accurately using as threshold (**Supplementary Fig. 8**). We corrected for *λ_GC_,* which was 1.723, for CEU&TSI. Adjustment for *λ_GC_* significantly reduced population stratification (**Supplementary Fig. 9**), and was consequently possible to filter out the baseline difference between these two cohorts. After correction, we found evidence of selection at the lactose persistence locus, *LCT* (*p*-value=1.21e-20). Due to hitchhiking effect, the region near *LCT* also showed divergent allele frequencies. For example, the *DARS* gene, 0.15M away from *LCT,* was also significantly associated with *E*_1_ (*p*-values=1.51e-23). *HERC2* was slightly below genome-wide significance level (*p*-value=8.22e-08), indicating that anthropological difference reflected geographic locations of two cohorts but not under selection as strong as *LCT.*

We then conducted EigenGWAS in the POPRES sample by treating *E*_1_ as a quantitative trait, and calculated the approximate *F_st_* for each SNP given two groups split by the threshold of *E*_1_ > 0 (**Supplementary Fig. 10**). Given 643,995 SNPs, the genome-wide threshold was *p*-value < 7.76e-08 for the significance level of *α* = 0.05. *λ_GC_* = 5.00, which indicated substantial population stratification as expected for POPRES. Correcting for *λ_GC_* systematically reduced the EigenGWAS *χ*^2^ test statistics (**Supplementary Fig. 11**), and we replicated the significance of *LCT* (*p*-value=1.23e-22) and *DARS* (*p*-value=8.99e-22) (**Table 2**), suggesting selection at these regions. *HERC2* was also replicated with *p*-value 8.15e-09, and with *F_st_* of 0.041.

**Table 2.** Gene discovery using EigenGWAS.

| Gene | Lead SNP | Position | Allele | $p$-value ($\lambda_{GC}$) | MAF (TSI:CEU) | $F_{st}$ | Annotation |
| --- | --- | --- | --- | --- | --- | --- | --- |
| CEU & TSI samples |  |  |  |  |  |  |  |
| **LCT** | rs6719488 | 2:135817629 | G/T | 6.68e-34 (1.21e-20) | 0.733:0.206 | 0.558 | Lactose persistent locus |
| **DARS** | rs13404551 | 2:135964425 | C/T | 8.18e-39 (1.51e-23) | 0.756:0.206 | 0.604 | Genetic hitchhiking due to *LCT*. |
| **MICA** | rs2256175 | 6:31412672 | T/C | 8.94e-10 (2.60e-6) | 0.665:0.360 | 0.183 | MHC class I polypeptide-related sequence A |
| **HIF1A** | rs2256205 | 14:61670944 | A/G | 1.51e-10 (8.86e-7) | 0.464:0.179 | 0.192 | HIF-1A thus plays an essential role in embryonic vascularization, tumor angiogenesis and pathophysiology of ischemic disease. |
| **HERC2** | rs8039195 | 15:26189679 | C/T | 2.75e-12 (8.22e-08) | 0.403:0.122 | 0.212 | Genetic variations in this gene are associated with skin/hair/eye pigmentation variability |
| POPRES European samples |  |  |  |  |  |  |  |
|  |  |  |  |  | Southern Europeans :<br>Northern Europeans |  |  |
| **LCT** | rs3754686 | 2:135817629 | T/C | 3.30e-106 (1.23e-22) | 0.514:0.279 | 0.110 |  |
| **DARS** | rs13404551 | 2:135964425 | C/T | 6.32e-102 (8.99e-22) | 0.518:0.293 | 0.106 |  |
| **SLC4A4** | rs605203 | 6:31819235 | C/A | 8.94e-44 (5.77e-10) | 0.214:0.343 | 0.040 | Defects in this gene can cause sialidosis, a lysosomal storage disease |
| **HERC2** | rs1667394 | 15:26189679 | C/T | 3.90e-38 (8.15e-09) | 0.276:0.173 | 0.041 |  |

Notes: The $p$ -value cutoff for CEU&TSI was 5.44e-08 (919,133 SNPs), for POPRES was 7.76e-08 (643,995 SNPs) at genome-wide significance level of $\alpha = 0.05$ . $\lambda_{GC} = 1.725$ for CEU&TSI, and $\lambda_{GC} = 5.00$ for POPRES.
$F_{st}$ is calculated by partitioning the sample into two groups upon $E_1 > 0$ . For TSI&CEU set, partitioning on $E_1$ perfectly separated TSI (88 samples) and CEU (112 samples). For POPRES, partitioning on $E_1$ separated southern European population (1,092 samples) and northern European population (1,374 samples).

### Prediction accuracy for projected eigenvector

We investigated three aspects of EigenGWAS prediction: 1) the number of loci needed to achieve high accuracy for the projected eigenvectors; 2) the required sample size of the training set; 3) the importance of matching the population structure between the training and the test sets.

Using the POPRES samples, we split 5% (125 individuals), 10% (250 individuals), 20% (500 individuals), 30% (750 individuals), 40% (1000 individuals), and 50% (1250 individuals) of the sample as the training set, and used the remainder of the samples as the test set. Eigenvectors were estimated using all markers in each training set. As predicted by our theory (Eq 7), the prediction accuracy of the projected eigenvector was consistent with 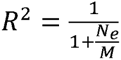 in which *N_e_* = 1,000 for *E*_1_ empirically. If only 100 and 1,000 random SNPs were sampled as predictors, the expected maximal *R*^2^ = 0.091 and 0.5, respectively and accuracy reached almost 1 if more than 100,000 SNPs were sampled. In agreement with our theory (**Fig. 6**), if the number of predictors were too small the prediction accuracy was poor, with prediction accuracy increasing with the addition of more markers for *E*_1_. When the sample size of the discovery was 1,000 or above, maximal prediction accuracy was achieved, as predicted in our theory. Therefore, a discovery with a sample size greater than 1,000 should be sufficient to predict the first eigenvector of an independent set, provided that population structure is the same across the discovery and prediction samples (**Fig. 6**). In contrast, the prediction accuracy for prediction eigenvectors decreased (**Fig. 6**) quickly for eigenvectors other than *E*_1_. For example, the prediction accuracy for *E*_2_ was below *R*^2^ < 0.2 and *R*^2^ < 0.15 for *E*_3_. For *E*_4_ ~ *E*_10_, the prediction accuracy dropped down to nearly zero. This is consistent with the top 2~3 eigenvectors explaining the majority of variation (McVean, 2009), if the training and the test sets had their population structure matched.

**Figure 6.**
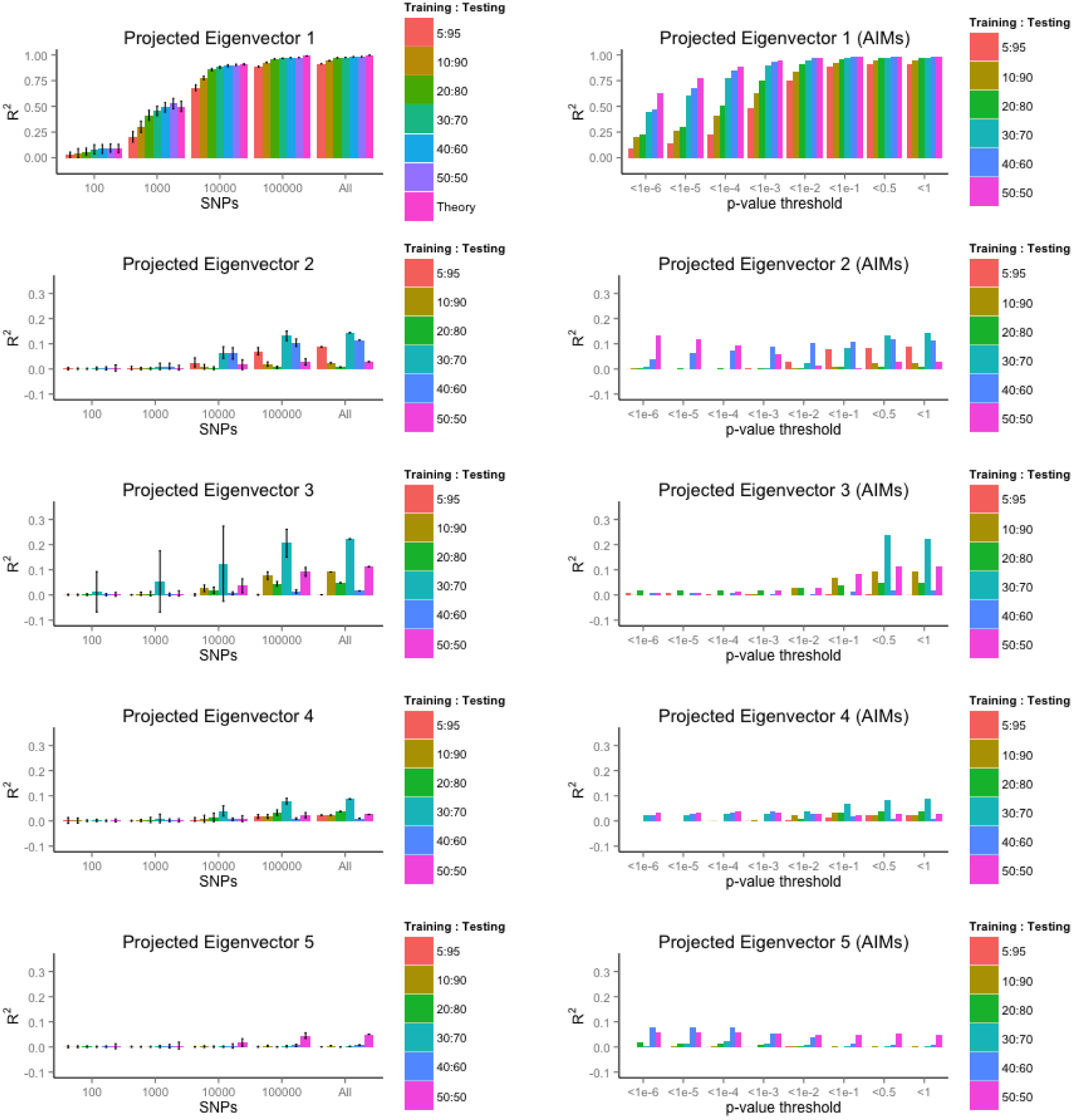
Prediction accuracy of the projected eigenvectors for POPRES samples. Given 2,466 POPRES samples, the data were split to 5%:95%, 10%:90%, 20%:80%, 30%:70%, 40%:60%, and 50%:50, as training and test set. The left columns represent prediction accuracy () using randomly selected numbers (100, 1,000, 10,000, 100,100, all) of markers, the 95% confidence interval were calculated from 30 replication for resampling given number of markers. In contrast, the right columns represent the predicted accuracy for 8 p-value thresholds (1e-6, 1e-5, 1e-4, 1e-3, 1e-2, 1e-1, 0.5, and 1) for EigenGWAS SNPs.

If EigenGWAS SNPs of low *p*-value were likely to be AIMs, we would hypothesise that AIM markers would be more efficient in giving high accuracy for the predicted eigenvectors (**Fig. 6**). For *E*_1_, the prediction accuracy reached 1 more quickly by using markers selected by *p*-value thresholds. The prediction accuracy for projected *E*_2_ was dependent upon the threshold. For projected *E*_2_ given a 50:50 split of POPRES sample, applying the threshold of *p*-value < 1e-6 (927 SNPs), *R*^2^ = 0.136, as high as using all markers. For other projected eigenvectors, the pattern of accuracy did not change much after applying *p*-value thresholds because in general, the prediction accuracy was low. This indicated that eigenvectors other than the first two eigenvectors capture little replicable population structure in POPRES.

In practice, the training and the test set may not match perfectly on population structure, and this will likely lead to a reduction in prediction accuracy. To demonstrate this, we split the POPRES samples into two sets: pooling Swiss (991 samples) and French (96 samples) samples into one group (SF), and the rest of the samples into the other group (NSF). We used SF as the training and the NSF as the testing. As SF was almost an average of North European and South European gene flow, making a less stratified population, its EigenGWAS effects would be consequently small and less “heritable”. When using all SNPs effects estimated from SF set, the observed prediction accuracy for NSF set was *R*^2^ = 0.33 and 0.005 for *E*_1_ and *E*_2_, respectively. These results indicate that a matched training and test set is important for prediction accuracy of the projected eigenvectors.

Ancestry information may still be elucidated well even if the training set and the test set do not match well in their population structure. Using HapMap3 as the training set, we also tried to infer the ancestry of the Puerto Rican cohort (PUR, 105 individuals) and Pakistani cohorts (PJL, 95 individuals) from 1000 Genomes project (The 1000 Genomes Project Consortium, 2012). In chromosome 1, 74,500 common SNs were found between HapMap3 and 1000 Genomes project. As illustrated, using only 74,500 common markers between HapMap3 and 1000 Genome projects SNPs on chromosome 1, it projected Eigenvectors accurately revealed the demographic history of Puerto Rican cohort, an admixture of African and European gene flows, and Pakistan cohort, an admixture of Asian and European gene flows (**Fig. 7**).

**Figure 7.**
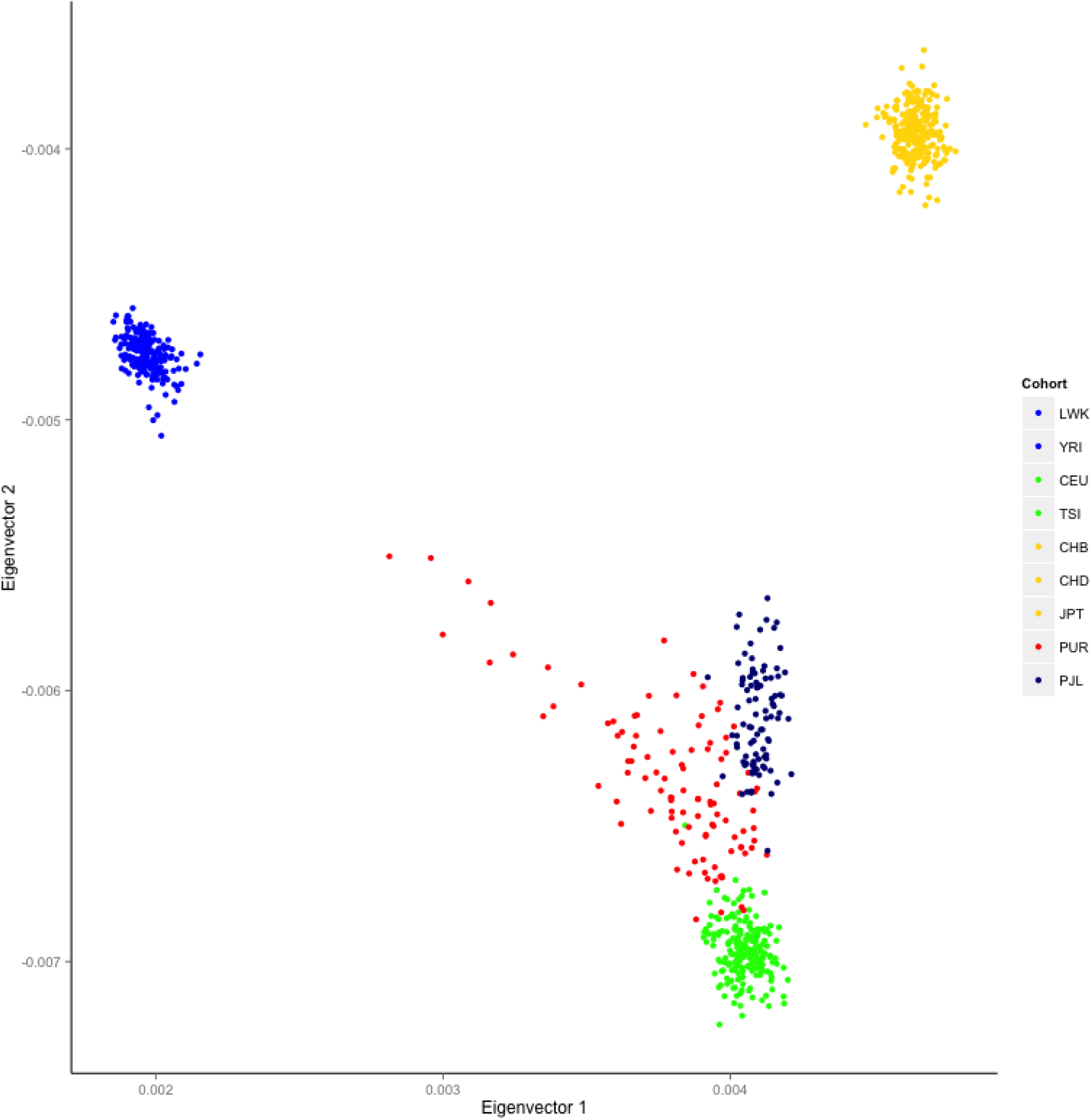
Projected eigenvectors for Puerdo Rican cohort (PUR) and Pakistan cohort (PJL) in 1000 Genome project. The training set was HapMap3 samples build on 919,133 SNPs. The eigenvectors 1 and 2 for were generated based on the 74,500 common SNPs on chromosome 1. PUR showed an admixture of African and European gene flows, and PJL Asian and European gene flows.

As a negative control, we replicated the prediction study for simulated data used in the previous section. The simulated data was split to two equal sample size. As there was no population structure in the simulated data, the prediction accuracy was poor, *R*^2^ = 0.01 from to *E*_1_ to *E*_10_. This demonstrates that prediction can be used to validate whether population structure exists within a genotype sample.

We concluded that to achieve high prediction accuracy of projected eigenvectors for independent samples, there are several conditions to be met: 1) the training set should harbour sufficient population stratification; 2) the sample size of the training should be sufficiently large; 3) the test sets should be as concordant as possible in its population structure; 4) when there is no real population structure, the prediction accuracy is very low close to zero; 5) depending on the population, high prediction was largely achievable for the projected *E*_1_.

## Discussion

Eigenvectors have been routinely employed in population genetics, and various approaches have been proposed to offer interpretation and efficient algorithms (Patterson *et al*., 2006; Rokhlin *et al.*, 2009; McVean, 2009; Chen *et al.*, 2013; Galinsky *et al.*, 2015). In this study, we created a GWAS framework for studying and validating population structure, and offer an interpretation of eigenvectors within this framework. The EigenGWAS framework (least square) identifies ancestry informative markers and loci under selection across gradients of ancestry.

We integrated SVD, BLUP, and single-marker regression into a unified framework for the estimation of SNP-level eigenvectors. SVD is a special case of BLUP when heritability is of 1 for the trait and the target phenotype is an eigenvector. Furthermore, the BLUP is equivalent to the commonly used GWAS method for estimating SNP effects. As demonstrated, the correlation between BLUP and GWAS is almost 1 for the estimated SNP effects. EigenGWAS offers an alternative way in estimating *F_st_* that can replace conventional *F_st_* when population labels are unknown, populations are admixed, or differentiation occurs across a gradient. As demonstrated for CEU&TSI samples, EigenGWAS brings out nearly identical estimation of *F_st_* compared with conventional estimation.

Different from conventional GWAS, which requires conventional phenotypes, the proposed EigenGWAS provides a novel method for finding loci under selection based on eigenvectors, which are generated from the genotype data itself. An EigenGWAS hit may reflect the consequence of process and thus additional evidence is needed to differentiate selection from drift. *LCT* is a known locus under selection, which differs in its allele frequency as indicated by *F_st_* statistic between Northern and Southern Europeans (Bersaglieri *et al*., 2004). We replicated the significance of *LCT* in CEU&TSI samples and POPRES European samples. *DARS* has been found in association with hypomyelination with brainstem and spinal cord involvement and leg spasticity (Taft *et al*., 2013). In addition, we also found *HERC2* locus independently, which may indicate the existence of anthropological difference in certain characters, such as hair, skin, or eyes color across European nations (Voight *et al*., 2006; Visser *et al.*, 2012).

Although by definition selection and genetic drift are different biological processes, both lead to allele frequency differentiation across populations and often difficult to tear them apart. In this study, with and without adjustment for *λ_GC_* from EigenGWAS offers a straightforward way to filter out population stratification. For example, with adjustment for *λ_GC_, LCT* and *DARS* were still significant in both EigenGWAS, while *HERC2* was only significant in POPRES. If adjustment for *λ_GC_* removed the average genetic drift since the most recent common ancestor for the whole sample, it might indicate that *HERC2* reflected the anthropological difference between subsamples but not under selection as strong as that for *LCT.* Nevertheless, *LCT* was differentiated due to selection that was on top of genetic drift, and for *DARS,* it might be significant due to hitchhiking effect. So, *LCT, DARS,* and *HERC2* were significant in EigenGWAS for different mechanisms.

In EigenGWAS application, it provides a clear scenario that *λ_GC_* is necessary if genetic drift/population stratification should be filtered out. It has been debated whether correction for *λ_GC_* is necessary for GWAS (Devlin and Risch, 1995; Yang, Weedon, *et al.*, 2011). If the inflation is due to population stratification, as initially introduced, it seems necessary to control for it. In contrast, if it is due to polygenic genetic architecture, then correction for *λ_GC_* will be a overkilling for GWAS signals. Interestingly, Patterson et al (Patterson *et al*., 2006) found that the top eigenvalues reflect population stratification, and in our study we found *λ_GC_* from EigenGWAS was numerically so similar to its corresponding eigenvalues. It in another aspect indicates *λ_GC_* captures population stratification. So, in concept and implementation, the correction for *λ_GC_* is technically reasonable. Of note, Galinsky et al also proposed a similar procedure to filter out population stratification in a study similar to ours (Galinsky *et al*., 2015), but we believe our framework is much easier to understand and implement in practice.

Once we have EigenGWAS SNP effects estimated, it is straightforward to project those effects onto an independent sample. The prediction of population structure was to that of recent studies (Chen *et al*., 2013). We found that the prediction accuracy for the top eigenvector could be as high as almost 1. Given a training set of about 1,000 samples, the prediction accuracy could be very high if there were a reasonable number of common markers in the order of 100,000. This number, which needs to be available in both reference set and the target set, is achievable. Further investigation may be needed to check whether this number of markers is related to effective number or markers after correction for linkage disequilibrium for GWAS data. When the population structure of the test sample resembles the training sample, high accuracy will be achieved for the leading projected eigenvectors. Therefore, this approach is likely to be extremely beneficial for extremely large samples, such as UK Biobank samples and 23andMe, both of which have more than half million samples where direct eigenvector analysis may be infeasible. Our results suggest that sampling about 1,000 individuals from the whole sample as the training set and subsequently project EigenGWAS SNP effects to the reminding samples will be sufficient to reach a reasonable high resolution of the population structure.

Many improvements to the inference of ancestry using projected eigenvectors have been suggested (Chen *et al*., 2013). As the concordance of population structure between the training and test sets is often unknown (population structure, upon from genetic or social-cultural perspectives, its definition can be difficult or controversial), improvement of the inference of ancestry may or may not be achieved dependent upon the scale of the precision required for a sample. However, for classification of samples at ethnicity level, projected eigenvectors are likely to have high accuracy, as demonstrated in the Puerto Rican cohort and the Pakistani cohort. Therefore, when identifying ethnic outliers, using projected eigenvectors from HapMap is likely to be sufficient in practice.

Eigenvector analysis of GWAS data is an important well utilized data technique, and here we show that its interpretation depends on many factors, such as proportion of different subpopulations, and *F_st_* between subpopulations. Our EigenGWAS approach provides intuitive interpretation of population structure, enabling ancestry informative markers (AIM) to be identified, and potentially loci under selection to be identified. To facilitate the use of projected eigenvectors, we provide estimated SNP effects from HapMap samples and POPRES and software that can largely reduce the logistics involved in conventional way in generating eigenvectors, such as reference allele match, and strand flips.

## Methods and Materials

**HapMap3 samples.** HapMap3 samples were collected globally to represent genetic diversity of human population (Altshuler *et al*., 2010). HapMap3 contains representative samples from many continents: CEU and TSI represent population from north and south Europe, CHB and JPT from East Asia, and CHD Chinese collected in Denver, Colorado. Loci with palindrome alleles (A/T alleles, or G/C alleles) were excluded, and 919,133 HapMap3 SNPs were used for the analysis.

**1000 Genomes project.** 1000 Genomes project samples were used as a prediction set for projecting eigenvectors (The 1000 Genomes Project Consortium, 2012). We selected the Puerto Rico cohort (PUR, 105 samples) and the Pakistan cohort (Punjabi from Lahore, Pakistan, 95 samples) for analysis.

**POPRES samples.** POPRES (Nelson *et al*., 2008) is a reference population for over 6,000 samples from Asian, African, and European nations. In this study, we selected 2,466 European descendants. The POPRES genotype sample was imputed to a 1000 Genomes reference panel (The 1000 Genomes Project Consortium, 2012). Imputation for the POPRES was performed in two stages. First, the target data was haplotyped using HAPI-UR (Williams *et al*., 2012). Second, Impute2 was used to impute the haplotypes to the 1000 genomes reference panel (Howie *et al*., 2011). We then selected SNPs which were present across all datasets at an imputation information score of >0.8. A full imputation procedure is described at https://github.com/CNSGenomics/impute-pipe. After quality control and removing loci with palindromic alleles (A/T alleles, or G/C alleles) 643,995 SNPs for POPRES remained. In addition, we also conducted the analysis using non-imputed 234,127 common markers between POPRES and HapMap3. As the results were between these two datasets were very similar, this report focused on the results from 643,995 SNPs, which were more informative.

**Simulation scheme I: null model without population structure.** 2,000 unrelated samples with 500,000 biallelic markers, which were in linkage equilibrium to each other, were simulated. The minor allele frequencies ranged from 0.01~0.5, and Hardy-Weinberg equilibrium was assumed for each locus. All individuals were simulated from a homogeneous population, with no population stratification. In order to calculate *F_st_* at each locus, we divided the sample into sub-populations based upon eigenvectors that were estimated from a genetic relationship matrix calculated using all 500,000 markers (see below).

**Simulation scheme II: null model with population structure.** In general, this simulation scheme was followed Price et al (Price *et al*., 2006). 2,000 unrelated samples with 10,000 biallelic markers, which were in linkage equilibrium to each other, were generated. For each marker, its ancestral allele frequency was sampled from a uniform distribution between 0.05 to 0.95, and its frequency in a subpopulation was sampled from Beta distribution with parameters 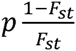 and 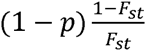. The Beta distribution had mean of *p* and sampling variance of *p*(1 − *p*)*F_st_.* Once the allele frequency for a subpopulation over a locus was determined as *p_s_*, individuals were generated from a binomial distribution *Binomil* (2, *p_s_*.). It agreed with the quantity that measures the genetic distance between a pair of subpopulations (Cavalli-Sforza *et al.*, 1996).

### Calculating individual-level eigenvectors

We assume that there is a reference sample consisting of *N* unrelated *M* individuals and markers. *X_i_* = (*X_i_*_1_*,X_i_*_2_*, …, X_iM_*)*^T^,* is a vector of the *i^th^* individual’s genotypes along *M* loci, with *x* the number of the reference alleles. An *N* × *N* genetic relatedness (correlation) matrix ***A*** (matrix in bold font) for each pair of individuals is defined as 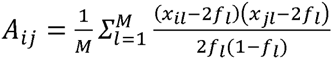, in which *f_l_* is the frequency of the reference allele. The principal component analysis (PCA) is then implemented on the *A* matrix (Price *et al*., 2006), generating 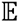, which is an *N* × *K* (*K ≤ N*) matrix, in which *E_k_* is the eigenvector corresponding to the *k^th^* largest eigenvector.

### Unified framework for BLUP, SVD, and EigenGWAS

Theoretically, PCA can also be implemented on a *M* × *M* matrix, but this is often infeasible because the *M* × *M* matrix is very large. However, for individual *i,* eigenvector *k* can also be written as:

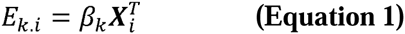

in which *β_k_* is a *M* × 1 SNP-level vector of the SNP effects on *E_k_,* and *x_i_* is the genotype of the *i^th^* individual across *M* loci. In the text below, we denote individual-level eigenvector as eigenvector *N* × 1 vector), and SNP-level eigenvector (*M* × 1) as SNP effects.

We review three possible methods to estimate *β* given eigenvectors. The first method is best linear prediction (BLUP), which is commonly used in animal breeding and recently has been introduced to human genetics for prediction (Henderson, 1975; Goddard *et al.*, 2009). The second method is to convert an individual-level eigenvector to SNP-level eigenvector using SVD, as proposed by Chen et al (Chen *et al*., 2013). The third method is the approach outlined here, EigenGWAS, which is a single-marker regression, as commonly used in GWAS analysis.

### Method 1 and 2: BLUP and SVD

For a quantitative trait, *y* = *μ* + *β****X*** + *e,* in which *y* is the phenotype, *μ* is the grand mean, *β* is the vector for additive effects, *X* is the genotype matrix, and *e* is the residual. Without loss of generality, the BLUP equation can be expressed as:

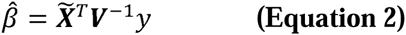

in which 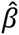 is the estimates of the SNP effects, 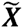 is the standardized genotype matrix, ***V*** is the variance covariance with 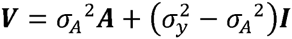, and is the trait of interest (Henderson, 1975). Replacing *y* with individual-level eigenvector (*E_k_*), Eq 2 can be written as

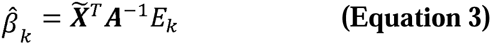

in which *β_k_* is the BLUP estimate of the SNP effects, *E_k_* is the *k^th^* eigenvector estimated from the reference sample., The ***V*** matrix can be replaced with ***A*** because the eigenvector has no residual error (i.e. *h*^2^=1). This method has also been proposed as an equivalent computing algorithm for genomic predictions (Maier *et al*., 2015).

In addition, the connection between PCA and SVD can be established through the transformation between the *N* × *N* matrix to the *M* × *M* matrix (McVean, 2009). Let ***A* = *PDP***^−1^ in which ***D*** is a *N* × *N* diagonal matrix with *λ_k_*, ***P*** is *N* × *N* matrix with the eigenvectors. ***B* = *X****^T^*(***PDP***^−1^)^−1^***P*** = ***X****^T^****PD****^−^*^1^, in which ***B*** is *M* × *M* matrix. This is equivalent to the equation used in Chen et al (Chen *et al*., 2013) where ***B****^T^* **= *D****^−^*^1^(***X****^T^****P***)*^T^*. Thus, eigenvector transformation can be viewed as a special case of BLUP in which the heritability is 1 (Eq 3). However, under SVD another analysis step is then required to evaluate the significance of the estimated SNP effect. In an EigenGWAS framework an empirical *p*-value is produced when estimating the regression coefficient.

### Method 3: estimating SNP effects on eigenvectors with EigenGWAS

Given the realized genetic relationship matrix *A,* for unrelated homogeneous (i.i.d.) samples, *E*(*A_ij_*) = 0 (*i* ≠ *j*), and consequently *E*(***A***) = ***I*, an identity matrix**. Due to sampling variance of the genetic relationship matrix ***A***, the off diagonal is a number slightly different from zero even for unrelated samples (Chen, 2014). If we replace the matrix with its mathematical expectation – the identity matrix, Equation 3 can be further reduced to 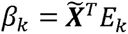, which is equivalent to single-marker regression *E_k_* = *a* + *bx* + *e*, as implemented in PLINK (Purcell *et al*., 2007). Furthermore, standardization for ***X*** is not required because it will not affect *p*-value. Thus, SNP effects can be estimated using the single-marker regression, which is computationally much easier in practice and is implemented in many software packages. Each SNP effect,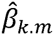, is estimated independently, and the *p*-value of each marker can be estimated, which requires additional steps in BLUP and SVD.

We summarise the properties and their transformation of SVD, BLUP, and EigenGWAS as below:

1. *E*_k_ is determined by the ***A*** matrix, or in another words, it is determined by the genotypes completely. If we consider each *E_k_* is the trait of interest – a quantitative trait, its heritability is 1.
2. *h*^2^ = 1. SVD and BLUP are both computational tool in converting a vector from *N* × *N* matrix to a *M* × *M* matrix. SVD is a special case to BLUP when *h*^2^ = 1 for BLUP.
3. *h*^2^ = 1 and *E*(***A***) = ***I***. When these two conditions are set, BLUP is further reduced to single-marker association studies, which is EigenGWAS as suggested in this study.

Recently, in an independent work Galinsky et al (Galinsky *et al*., 2015) introduced an approximation to find the proper scaling for SNP effects (“SNP weight” in Galinsky’s terminology) estimated from SVD, in order to produce accurate *p*-values. In our EigenGWAS framework, *p*-values for individual-level SNP eigenvector are automatically generated. In practice, it is conceptually easier to conduct EigenGWAS on eigenvectors than to conduct BLUP/SVD. Also, if computational speed is of concern, EigenGWAS can be easily parallelized for each chromosome, each region, or even each locus.

### Interpretation for EigenGWAS

We can write a linear regression model *E_k_* = *a* + *βx* + *e*, in which both *E_k_* and *x* is standardized. Assuming that a sample has two subdivisions, which have sample size *n*_1_ and 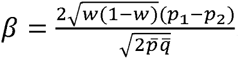, and the sampling variance for *β* is 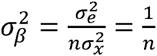. A chi-square test for *β* is

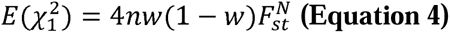

in which 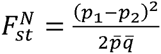 is Nei’s estimator of genetic difference for a biallelic locus (Nei, 1973).

In principal component analysis, the proportion of the variance explained by the largest eigenvalue is equal to 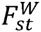 (McVean, 2009), in which 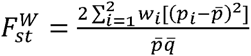 for a pair of subpopulations as defined in Weir (Weir, 1996). So 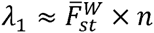, in which 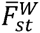 characterizes the average divergence for a pair of subpopulations. When the test statistic, Eq 4, is adjusted by the largest eigenvalue *λ*_1_, an equivalent technique in GWAS for the correction of population stratification, 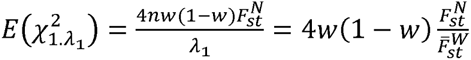. For a population with a pair of subdivisions 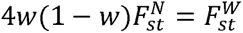. So

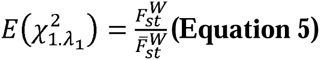

after the adjustment of the largest eigenvalue, the test statistic immunes of population stratification, at least for a divergent sample.

For a locus under selection, which should have a greater *F_st_* than 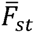 the background divergence. So the statistical power for detecting whether a locus is under selection is determined by the strength of selection, which can be defined as the ratio between *F_st_* of a particular locus and 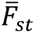 the average divergent in the sample. It is analogous to consider a chi-square test with non-centrality parameter (NCP), 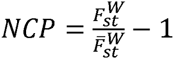.

Otherwise specified, in this study *F_st_* is referred to the one defined in Weir (Weir, 1996).

### Validation and prediction for population structure

Once *β_k_* is estimated, it is straightforward to get genealogical profile for an independent target sample. In general, it is equivalent to genomic prediction, and the theory for prediction can be applied (Daetwyler *et al*., 2008; Dudbridge, 2013). The predicted genealogical score can be generated as

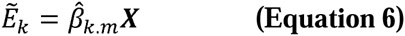

in which *E_k_* is the predicted *k^th^* eigenvector, 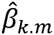 is the estimated SNP effects, and ***X*** is the genotype for the target sample. We focus on the correlation between the predicted eigenvectors and the direct eigenvectors, and thus it does not matter whether ***X*** or 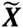 is used.

In contrast to conventional prediction studies, which focus on a metric phenotype of interest, prediction of population structure is focussed on a “latent” variable. This latent variable is the genetic structure of population, which is shaped by allele frequency and linkage disequilibrium of markers. Thus, expectations of prediction accuracy differ from what has been established for conventional prediction (Daetwyler *et al*., 2008; Dudbridge, 2013) 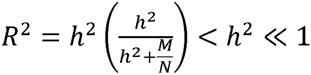. We therefore assess prediction of accuracy for *E*_1_ across markers, when using different prediction thresholding (Purcell *et al*., 2009).

Here we proposed an equation for prediction accuracy, especially for *E*_1_

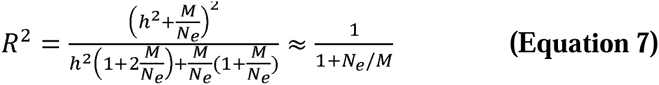

when there is no heritability, the predictor can be simplified to 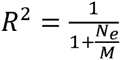, meaning that as the number of markers increases prediction accuracy should rapidly reach 1. Here the *h*^2^ is interpreted as the genetic difference in the source population, or real ancestry informative markers. For a homogeneous population, the genetic difference is large due to genetic drift, and *h*^2^ ≈ 0.

For this study, the genetic relationship matrix (***A*** matrix), principal component analysis, and BLUP estimation were conducted using GCTA software (Yang, Lee, *et al.*, 2011). Single-marker GWAS was conducted using PLINK (Purcell *et al*., 2007), or GEAR (https://github.com/gc5k/GEAR/wiki/EigenGWAS; https://github.com/gc5k/GEAR/wiki/ProPC).

## Web resource and data availability

GEAR is available at http://cnsgenomics.com/

GCTA is available at http://cnsgenomics.com/

PLINK is available at http://pngu.mgh.harvard.edu/~purcell/plink/index.shtml 1000 Genomes Project: http://www.1000genomes.org/

## Acknowledgements

This research was funded by ARC (DE130100614 to SHL), NHMRC (APP1080157 to SHL, APP1084417 and APP1079583 to BB, and APP1050218 to MRR), and GBC was supported by IAP P7/43-BeMGI from the Belgian Science Policy Office Interuniversity Attraction Poles (BELSPO-IAP) program. We thank Peter M. Visscher for discussion, helpful comments, and for proposing the name EigenGWAS. Robert Maier assisted with ggplot, and Alex Holloway helped with Github. We also thank to the Information Technology group, the Queensland Brain Institute. The POPRES dataset were obtained from dbGaP at http://www.ncbi.nlm.nih.gov/gap through accession number phs000145.v4.p2.

## Author contributions

GBC, SHL, and BB conceived study. GBC, SHL, BB, and MRR designed the experiment. GBC and SHL developed the theory and methods. BB conducted the quality control for HapMap data, and MRR conducted quality control for POPRES data. GBC performed the analyses of the study. GBC and ZXZ developed GEAR software. GBC, MRR, SHL, and BB wrote the paper.

## References

Altshuler DM, Gibbs RA, Peltonen L, Dermitzakis E, Schaffner SF, Yu F, et al. (2010). Integrating common and rare genetic variation in diverse human populations. Nature 467: 52–8.

Bersaglieri T, Sabeti PC, Patterson N, Vanderploeg T, Schaffner SF, Drake JA, et al. (2004). Genetic Signatures of Strong Recent Positive Selection at the Lactase Gene. Am J Hum Genet 74: 1111–1120.

Bryc K, Bryc W, Silverstein JW (2013). Separation of the largest eigenvalues in eigenanalysis of genotype data from discrete subpopulations. Theor Popul Biol 89: 3443.

Cavalli-Sforza LL, Menozzi P, Piazza A (1996). The History and Geography of Human Genes. Princeton University Press.

Chen G-B (2014). Estimating heritability of complex traits from genome-wide association studies using IBS-based Haseman-Elston regression. Front Genet 5: 107.

Chen C-Y, Pollack S, Hunter DJ, Hirschhorn JN, Kraft P, Price AL (2013). Improved ancestry inference using weights from external reference panels. Bioinformatics 29: 1399–406.

Daetwyler HD, Villanueva B, Woolliams J a (2008). Accuracy of predicting the genetic risk of disease using a genome-wide approach. PLoS One 3: e3395.

Devlin B, Risch N (1995). A comparison of linkage disequilibrium measures for fine-scale mapping. Genomics 29: 311–22.

Devlin B, Roeder K (1999). Genomic control for association studies. Biometrics 55: 997–1004.

Dudbridge F (2013). Power and predictive accuracy of polygenic risk scores. PLOS Genet 9: e1003348.

Galinsky KJ, Bhatia G, Loh P, Georgiev S, Mukherjee S, Nick J (2015). Fast principal components analysis reveals independent evolution of ADH1B gene in Europe and East Asia. bioRxiv. http://dx.doi.org/10.1101/018143.

Goddard ME, Wray NR, Verbyla K, Visscher PM (2009). Estimating Effects and Making Predictions from Genome-Wide Marker Data. Stat Sci 24: 517–529.

Henderson CR (1975). Best Linear Unbiased Estimation and Prediction under a Selection Model. Biometrics 31: 423–447.

Howie B, Marchini J, Stephens M, Chakravarti A (2011). Genotype Imputation with Thousands of Genomes. G3 1: 457–470.

Maier R, Moser G, Chen G, Ripke S, Group CW, Consortium PG, et al. (2015). Joint Analysis of Psychiatric Disorders Increases Accuracy of Risk Prediction for Schizophrenia, Bipolar Disorder, and Major Depressive Disorder. Am J Hum Genet 96: 283–294.

McVean G (2009). A genealogical interpretation of principal components analysis. PLOS Genet 5: e1000686.

Nei M (1973). Analysis of gene diversity in subdivided populations. Proc Natl Acad Sci U S A 70:3321–3.

Nelson MR, Bryc K, King KS, Indap A, Boyko AR, Novembre J, et al. (2008). The Population Reference Sample, POPRES: A Resource for Population, Disease, and Pharmacological Genetics Research. Am J Hum Genet 83: 347–358.

Novembre J, Johnson T, Bryc K, Kutalik Z, Boyko AR, Auton A, et al. (2008). Genes mirror geography within Europe. Nature 456: 98–101.

Patterson N, Price AL, Reich D (2006). Population structure and eigenanalysis. PLOS Genet 2: e190.

Price AL, Patterson NJ, Plenge RM, Weinblatt ME, Shadick N a, Reich D (2006). Principal components analysis corrects for stratification in genome-wide association studies. Nat Genet 38: 904–9.

Purcell S, Neale B, Todd-Brown K, Thomas L, Ferreira M a R, Bender D, et al. (2007). PLINK: a tool set for whole-genome association and population-based linkage analyses. Am J Hum Genet 81: 559–75.

Purcell SM, Wray NR, Stone JL, Visscher PM, O’Donovan MC, Sullivan PF, et al. (2009). Common polygenic variation contributes to risk of schizophrenia and bipolar disorder. Nature 460: 748–752.

Reich D, Thangaraj K, Patterson N, Price AL, Singh L (2009). Reconstructing Indian population history. Nature 461: 489–94.

Rokhlin V, Szlam A, Tygert M (2009). A randomized algorithm for principal component analysis. SIAM J Matrix Anal Appl 31: 1100–1124.

Taft RJ, Vanderver A, Leventer RJ, Damiani S a, Simons C, Grimmond SM, et al. (2013). Mutations in DARS cause hypomyelination with brain stem and spinal cord involvement and leg spasticity. Am J Hum Genet 92: 774–80.

The 1000 Genomes Project Consortium (2012). An integrated map of genetic variation from 1,092 human genomes. Nature 491: 56–65.

Visser M, Kayser M, Palstra R-J (2012). HERC2 rs12913832 modulates human pigmentation by attenuating chromatin-loop formation between a long-range enhancer and the OCA2 promoter. Genome Res 22: 446–55.

Voight BF, Kudaravalli S, Wen X, Pritchard JK (2006). A map of recent positive selection in the human genome. PLOS Biol 4: e72.

Weir BS (1996). Genetic data analysis, 2nd edn. Sinauer Associates, Inc.: Sunderland, MA, USA.

Williams AL, Patterson N, Glessner J, Hakonarson H, Reich D (2012). Phasing of many thousands of genotyped samples. Am J Hum Genet 91: 238–251.

Yang J, Lee SH, Goddard ME, Visscher PM (2011). GCTA: a tool for genome-wide complex trait analysis. Am J Hum Genet 88: 76–82.

Yang J, Weedon MN, Purcell S, Lettre G, Estrada K, Willer CJ, et al. (2011). Genomic inflation factors under polygenic inheritance. Eur J Hum Genet 19: 807–12.

Zhu X, Li S, Cooper RS, Elston RC (2008). A Unified Association Analysis Approach for Family and Unrelated Samples Correcting for Stratification. Am J Hum Genet 82: 352–365.

